# Barley disease screening: a multiplex digital droplet PCR approach for the detection of *Ramularia collo-cygni, Rhynchosporium graminicola* and *Pyrenophora teres*

**DOI:** 10.1101/2024.12.18.629171

**Authors:** Rabisa Zia, Zoë A. Popper, Steven Kildea

## Abstract

Barley is the most widely grown cereal crop in Ireland and faces substantial yield loss annually due to airborne, seed-borne, and soil-borne pathogens. The most prevalent and destructive of these include ramularia leaf spot, barley leaf scald and net blotch, caused by *Ramularia collo-cygni, Rhynchosporium graminicola* and *Pyrenophora teres* respectively. Infected seeds are considered as the major source of inoculum for these diseases. The absence of varietal resistance for these fungal pathogens suggests the need for better disease management strategies. Accurate pathogen detection is one of the most widely accepted disease management strategies. To facilitate such detections in complex samples, such as seed, we demonstrate two digital droplet PCR (ddPCR) assays, with the initial assay capable of detecting *R. graminicola, R*.*collo-cygni* and *P. teres*. For those identified as positive for *P. teres* samples, a form-specific assay was designed to distinguish between the two economically significant forms of *P. teres*; *P. teres f. maculata* (Ptm; spot form of net blotch) and *P. teres f. teres* (Ptt; net form of net blotch). The assay offers sensitive and specific pathogen detection from barley seeds up to pico moles/µl level. Limits of detection for *R. graminicola, R. collo-cygni* and *P. teres* in the triplex assay were calculated as 1.67 cp/µl, 1.06 cp/µl and 4.21 cp/µl respectively. For form-specific assay, limits of detection were calculated as 1.44 cp/µl for Ptt and 0.82 cp/µl for Ptm assay. Finally, the efficacy of the assay was demonstrated by screening a collection of historical barley seeds.

## 1 INTRODUCTION

After wheat and corn, barley is the 3^rd^ most widely produced cereal crop in Europe, making up almost 20% of the total cereals produced (European Union Crop Production, 2024). Due to the combination of agronomic and climatic factors, the potential for high yields and favourable end-markets, barley accounts for approximately 70% of all cereals produced in Ireland (CSO, 2023). Unfortunately, as with all crops, barley can succumb to a spectrum of diseases that if left unchecked can seriously impact potential yields, both quantitatively and qualitatively. Most notable amongst these are the fungal foliar diseases ramularia leaf spot (RLS) caused by *Ramularia collo-cygni*, barley scald caused by *Rhynchosporium graminicola*, and the spot and net forms of net blotch (SFNB and NFNB), caused by *Pyrenophora teres f. maculata* (Ptm) and *P. teres f. teres* (Ptt), respectively (Huss et al. 1992;Avrova & Knogge, 2012; Liu et al., 2011).

As is the case with all plant diseases and pests, limiting initial inoculum/pest sources is critical to achieving sustainable control and is the principal tactic of integrated pest management (Barzman et al. 2015). For *R. collo-cygni, R. graminicola* and *P. teres* trash and stubble from the previous season’s crops, volunteers and infected seed, all act as potential sources of initial inoculum. For *R. graminciola* and *P. teres*, it has been previously demonstrated that seed-borne infection represents a significant means of disease transmission and can directly impact disease levels and subsequent impacts on barley yields (Fountaine et al. 2010; Hysing & Wiik, 2013). For *R. collo-cygni*, whilst the pathogen can be readily detected in seed, and it has been demonstrated that it can grow from the seed into the developing seedling, the importance of seed-borne infection in the aetiology of RLS remains to be determined (Havis et al. 2014).

Traditionally seed has been tested for the presence of pathogens using culture dependent techniques, whereby the test seeds are cultured on a selective growth media and subsequent microbial growth is examined morphologically and/or microscopically to confirm the contaminating species (Hiddink et al. 2023). However, there is a major disadvantage of culture-dependent assays which is that they are restricted to those pathogens that can be easily grown in culture and have a relatively fast growth rate. Of the three major barley pathogens identified above, it is only possible to routinely identify *P. teres* through culture-dependent assays as both *R. graminicola* and *R. collo-cygni* are relatively slow growing and are outcompeted by other fungi (pathogenic or non-pathogenic) present in barley seed. Identification of *P. teres* in culture can also be difficult as it can be mistaken for *P. graminea*, which exhibits a similar morphology in culture (Liu et al. 2011). Similarly, it is not possible to distinguish between Ptm and Ptt when grown in culture or by using osmotic or blotter tests (Brodal, 1997). More recently developed detection methods that rely on detection of nucleic acids (either DNA or RNA) have improved specificity compared to culture-based detection methods. They also allow a wider range of pathogens to be detected, whilst simultaneously reducing costs and run time (Elnifro Elfath et al. 2000).

Seed samples are unfortunately often complex matrixes, which can inhibit or impair DNA based reactions and assay sensitivities (Lu et al. 2020), in particular those such as quantitative polymerase chain reaction (qPCR) on which quantification is based on comparable efficiencies of reactions required between the test sample and samples/standards of a known DNA concentration (Hindson et al. 2013). Digital droplet PCR (ddPCR, also referred to as digital PCR (dPCR) or crystal digital PCR (cdPCR) depending on the platform used) is a third generation of PCR in which the starting DNA sample is partitioned through the use of microfluidics into thousands of samples which subsequently undergo PCR reactions. By coupling partitioning and PCR with fluorescent dye based DNA probes it is possible to use endpoint analysis to confirm the presence or absence of the targeted DNA in partitioned samples (Mao et al. 2019). Through the use of Poisson’s Law, which estimates the proportion of droplets containing the target DNA, it is then possible to determine the total copies of the target DNA in the starting sample (Quan et al. 2018). As quantification is based on an end point reaction ddPCR is not sensitive to PCR inhibitors while qPCR is strongly influenced by the efficiencies of the reaction (Basu, 2017). Therefore, ddPCR allows for the accurate and sensitive quantification of specific DNA targets, even in complex matrices, making it an ideal platform for the detection of pathogens in cereal seeds.

Different PCR based methods have been developed for each of the commonly occurring barley pathogens, with qPCR assays available for both *P. teres* and *R. graminicola* (Leisova et al. 2006); Fountaine et al. 2007), and a dPCR based method for detection of *R. collo-cygni* recently published by Knight et al. (2022). To exploit the multi-spectral capacity of the Stilla 3-colour Nacia® ddPCR system, which allows simultaneous detection of the fluorescence emitted from three spectrally different fluorophores, here we combine the *R. graminicola* and *R. collo-cygni* assays of Fountaine et al. (2007) and Knight et al. (2022), with a *P. teres* probe based assay to provide a triplex ddPCR assay to simultaneously detect and quantify the presence of the three pathogens from historical barley seed samples. For the samples positive for *P. teres*, a further form-specific ddPCR assay was designed to detect and differentiate between the spot and net form of *P. teres*.

## 2 MATERIALS AND METHODS

A multistage process of assay design, optimisation and application was developed and is outlined in Figure 1.

**Figure 1.**
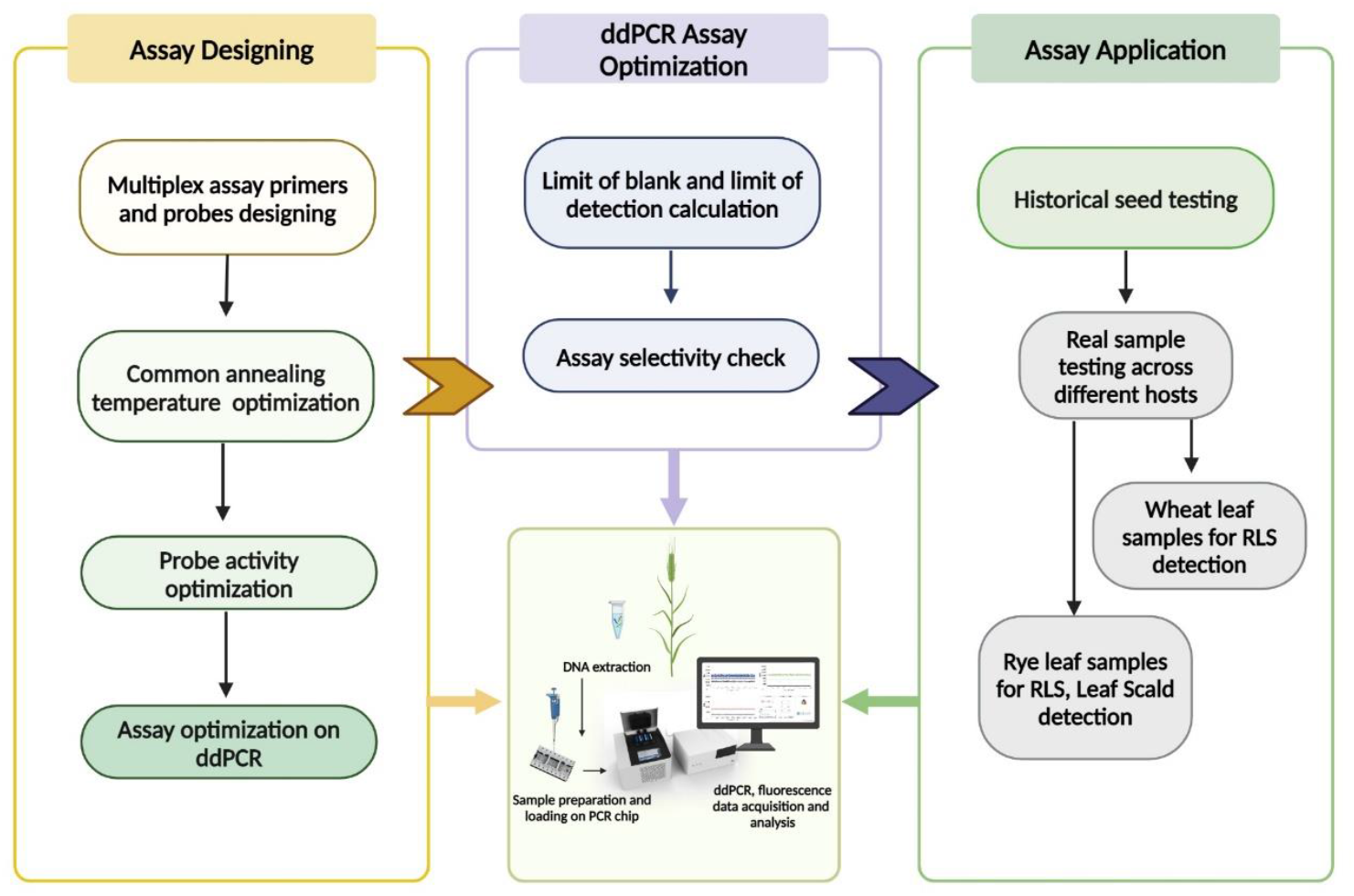
Summarized workflow of the digital droplet PCR assay (Created with BioRender.com)

### 2.1 Fungal strains and DNA extraction

All fungal cultures used in the study (Supplementary Table 1) were acquired from the phytopathological collection in Teagasc, Oak Park, Co. Carlow. DNA was extracted from these cultures following freeze drying (for 24 hours) and homogenisation (with Retsch Mixer Mill MM 400 at a frequency of 20 Hz for 5 minutes) using a Qiagen DNeasy Plant Mini kit (Qiagen, Germany), following the manufacturer’s instructions. Extracted DNA was quantified using Invitrogen™ Qubit™ Fluorometer with a Qubit™ 1X dsDNA BR Assay (Invitrogen, US). For some of the cultures (See Supplementary Table 1: Isolates utilized in the study), only DNA was provided and the depositor confirmed presence of fungal DNA with PCR.

### 2.2 Assay designs and initial optimization

For the initial ddPCR triplex assay to detect and distinguish between the three target fungal pathogens, the published fluorescence probe based assays for the detection of *R. collo-cygni* (Knight et al. 2022) and *R. secalis* (Fountaine et al. 2007) were selected. The *R. secalis* probe length was one base pair shorter than the original probe described by Fountaine et al. (2007). A separate set of primers and probe to detect *P. teres* was designed using Primer 3Plus (https://www.primer3plus.com/index.html) based on the Internal Transcribed Spacer (ITS) region of *P. teres* (OM471876.1). To further distinguish between Ptm and Ptt primers were deployed from Poudel et al. (2017) while probes were designed using Ptm and Ptt mRNA sequences from the same (accession numbers Ptm: KX909562.1 and Ptt: KX909556.1). However, for Ptm detection, a new reverse primer was designed using the same sequence in order to reduce the amplicon length for optimal qPCR efficiency (Van Holm et al. 2021) Online tool; PrimerQuest™ (Integrated DNA Technologies) was deployed for primer and probe designing. tripl Primers and probes were ordered from Integrated DNA Technologies (Europe). Details of all primers and probes are summarized in Table 1. Initial primer and probe optimisation, including identifying annealing temperatures, was conducted using routine PCR and visualised using agarose gel electrophoresis.

**Table 1.**
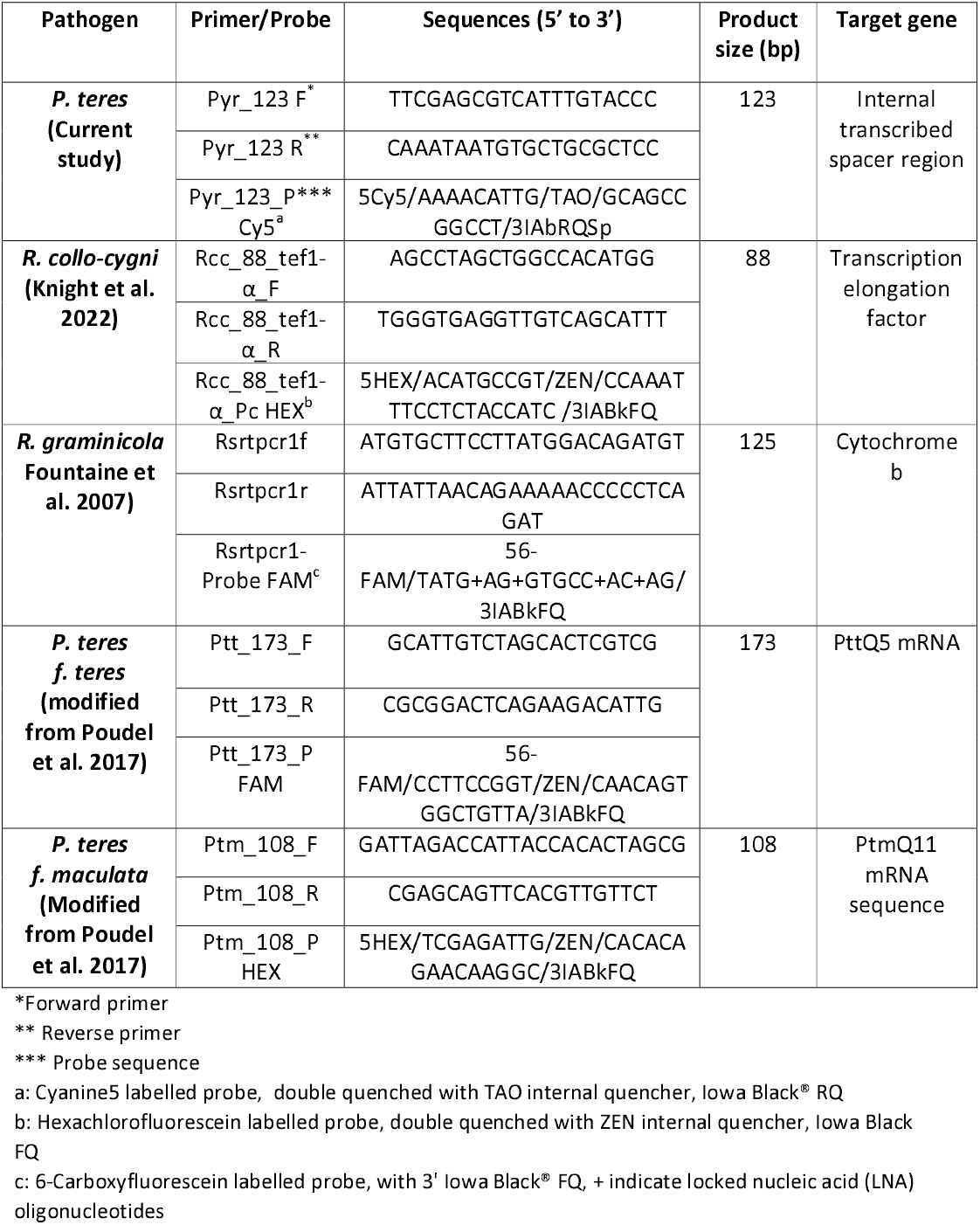
List of primers and probes deployed in this study.

### 2.3 ddPCR optimisation

Following these initial validation steps, the assays were transferred to ddPCR system (naica® Crystal Digital PCR 3 colour system, Stilla Technologies, France). ddPCR using the naica® Crystal Digital PCR 3 colour is separated into four stages: 1) the reaction mixture (primers, probes, DNA of the pathogen and host and ultrapure water) is partitioned into nano-sized droplets on a chip (up to 17k on a Ruby chip or 30k on a Sapphire chip), 2) partitioned samples undergo up to 40 PCR cycles, 3) once the PCR is complete a scan of the chip is taken under different wavelengths (corresponding to the excitation of the FAM, HEX and Cy5 dyes) is undertaken, and 4) scanned chips are analysed to identify droplets with the target DNA amplified based on presence or absence of the specified molecular probe-associated fluorescence. In the present study all reactions were conducted using the Ruby Chip (up-to 17k droplets per chamber) and 5X naica® multiplex PCR MIX (Stilla Technologies, France) and only reactions with >10k analysable droplets were included.

An initial series of reactions were performed to optimize concentrations of primers and probes required to differentiate between positive and negative droplets. Reaction mixtures were prepared in accordance with the guidelines provided by Stilla Technologies, consisting of 1 µl of Buffer A, 0.2 µl of Buffer B (both Buffer A and Buffer B were supplied as part of the 5X naica® multiplex PCR MIX), 0.25 µl each of the primers (0.05 µM final concentration), 0.15 µl of each probe (0.06 µM final concentration), 0.85 µl of barley DNA (17 ng/µl), 1 µl of original pathogen DNA was used as a template (0.1 ng/µl) and nuclease free water (Fisher Scientific, UK) was added to reach a final volume of 5 µl. All PCR reactions were performed in triplicates and in the presence of pathogen-free barley DNA as a matrix; extracted from two-week barley seedlings (variety: RGT Planet, Goldcrop Ltd, Ireland) grown under controlled conditions (16 h of light, 8 h of dark, at 20 and 16 °C). Thermal cycling conditions for both assays are as follows; partition at 25°C followed by 95°C for 3 minutes, then 5 cycles of 95°C for 10 seconds followed by 66°C for 30 seconds, and then 35 cycles at 95°C for 10 second and 64°C for 15 second. A compensation matrix was designed according to the guidelines provided by Stilla Technologies, to normalize the fluorescence overlap and spill over.

### 2.4 Determining Limits of Blank (LoB) and Limits of Detection (LoD)

Limit of blank (LoB) and Limit of detection (LoD) for the assays were calculated in multiplex fashion and in accordance with the guidelines provided by Stilla Technologies. LoB can be defined as the maximum target concentration expected in the blank sample with a probability PLoB = 1 – α (typically 1 – α = 95% where α = 5% of false positives), where α is the probability of making a false positive decision in a real blank sample (no DNA of any target except the background/barley DNA). Whereas, LoD is defined as the minimum concentration, a non-zero, statistically significant value, that can be detected with a probability of 1 – β (typically 1 – β = 95% where β = 5% of false negatives), β is the probability making a false negative decision in a real positive sample.

To determine the LoB, 30 blank (negative for the target for which LoB is being performed) reaction replications were performed for each target. To calculate LoD, ten-fold serial dilutions of a known target DNA concentration was performed and a non-zero, statistically significant concentration detected by the assay, designated as low-level sample, was calculated. Afterwards, five independent low-level samples were prepared and six reaction replications of each were performed. Based on the copy per microliter (cp/µl) measured by the fluorescence detector, LoD and LoB was calculated by using the online tool provided by Stilla Technologies (https://www.stillatechnologies.com/digital-pcr/statistical-tools/limit-detection/).

### 2.5 Linear dynamic range of the assay

To calculate the relationship between the expected and measured quantities of each of the target pathogen DNA, linear dynamic range of the assay was calculated. To achieve this, a 10-fold dilution series (0.01–0.00001 ng/µl) of each pathogen DNA (*R. collo-cygni, R. graminicola* and *P. teres*) was prepared and tested individually whilst keeping a constant DNA concentration of the other two pathogens (0.1 ng/µl) in the reaction mixture. The log10 values of concentration against copy number/µl (cp/ µl) were plotted and correlation was calculated in Microsoft Excel.

### 2.6 Specificity of the assays

Specificity of triplex and form-specific assays was performed by challenging the system with a number of barley pathogens/barley microflora including *Botrytis cinerea (Bc), Parastagonospora nodorum (Pn), Alternaria alternate (Aa), Fusarium graminearum (Fg), Microdochium nivale (Mn)* and *Fusarium culmorum (Fc)* accessions that are stored as part of the Phytopathological collection in Teagasc, Oak Park, Co. Carlow. Additionally, specificity of the form-specific was also verified against *R. collo-cygni* and *R. graminicola* in a singleplex reaction. Each of the pathogen DNA was tested in an individual ddPCR reaction.

### 2.6 Detection of pathogens in seed and leaf samples

To validate the reliability of the assays (triplex and form-specific assay) for naturally infected barley samples, historical winter (35 samples) and spring (32 samples) barley seed DNA from 2016–2018, with variable susceptibility to leaf blotches, were evaluated using the assays. The details of seed DNA extraction is described in Mulhare et al. (2021). Any droplet population above LoD of the respective assay was considered positive. Samples positive for *P. teres* were further tested by the form specific assay to confirm *P. teres* form.

Volunteers from the previous season crop are one of the sources of inoculum for RLS, barley leaf scald and net blotch. Efficacy of the triplex assay was also assessed in wheat volunteers from barley field with visual symptoms of RLS (Teagasc wheat trials in Knockbeg, Ireland (2023)). Leaves were freeze dried and DNA was extracted as described above. Extracted DNA (1 µl) was used as a template in the reaction mixture described in section 2.3.

## 3 Results

### 3.1 Assay design and optimization

For the detection of *R. collo-cygni* and *R. graminicola*, primers and probes were deployed from published assays targeting translation elongation factor 1-alpha gene and Cytochrome b genes respectively (Fountaine et al. 2007; Knight et al. 2022). However, for *P. teres* detection and to distinguish between both Ptt and Ptm it was necessary to design novel fluorescent probes (Table 1). The initial triplex assay was specific for the three pathogens, with no fluorescence overlap observed between the different probes-pathogen combinations (Figure 2A). In triplex assay, we observed 5044 cp/µl and 10210 cp/µl for *R. graminicola* and *P. teres*, for *R. collo-cygni and P. teres* 156.4 and 10993 cp/µl and for *R. graminicola, R. collo-cygni* sample 5600 and 164.4 cp/µl respectively. *teres* form specific assay there was no fluorescence overlap between the probe-pathogen combination, demonstrating specificity of the assay. We observed 112.8 cp/µl in FAM channel and 7086 cp/µl in Cy5 channel in Ptt sample, 576.6 cp/µl in HEX and 20267 cp/µl in Cy5 channel in Ptm sample, for both forms we observed 186.5 cp/µl in FAM, 501.3 cp/µl in HEX and 26113 cp/µl in Cy5 channel (Figure 2B). For *R. graminicola, R. collo-cygni and P. teres* the LoB were calculated to be 0.68, 0, and 1.34 cp/µl respectively. For the *P. teres* form-specific assay and LoB of 0 cp/µl was calculated for both Ptt and Ptm. Limits of detection (LoD) for *R. graminicola, R. collo-cygni* and *P. teres* in the triplex assay were calculated as 1.67 cp/µl, 1.06 cp/µl and 4.21 cp/µl respectively. Additionally, LoD was calculated as 1.44 cp/µl for Ptt assay and 0.82 cp/µl for Ptm assay. No positive droplets, irrespective of probe, were detected in negative control (barley DNA only) (Figure 2A & B).

**Figure 2.**
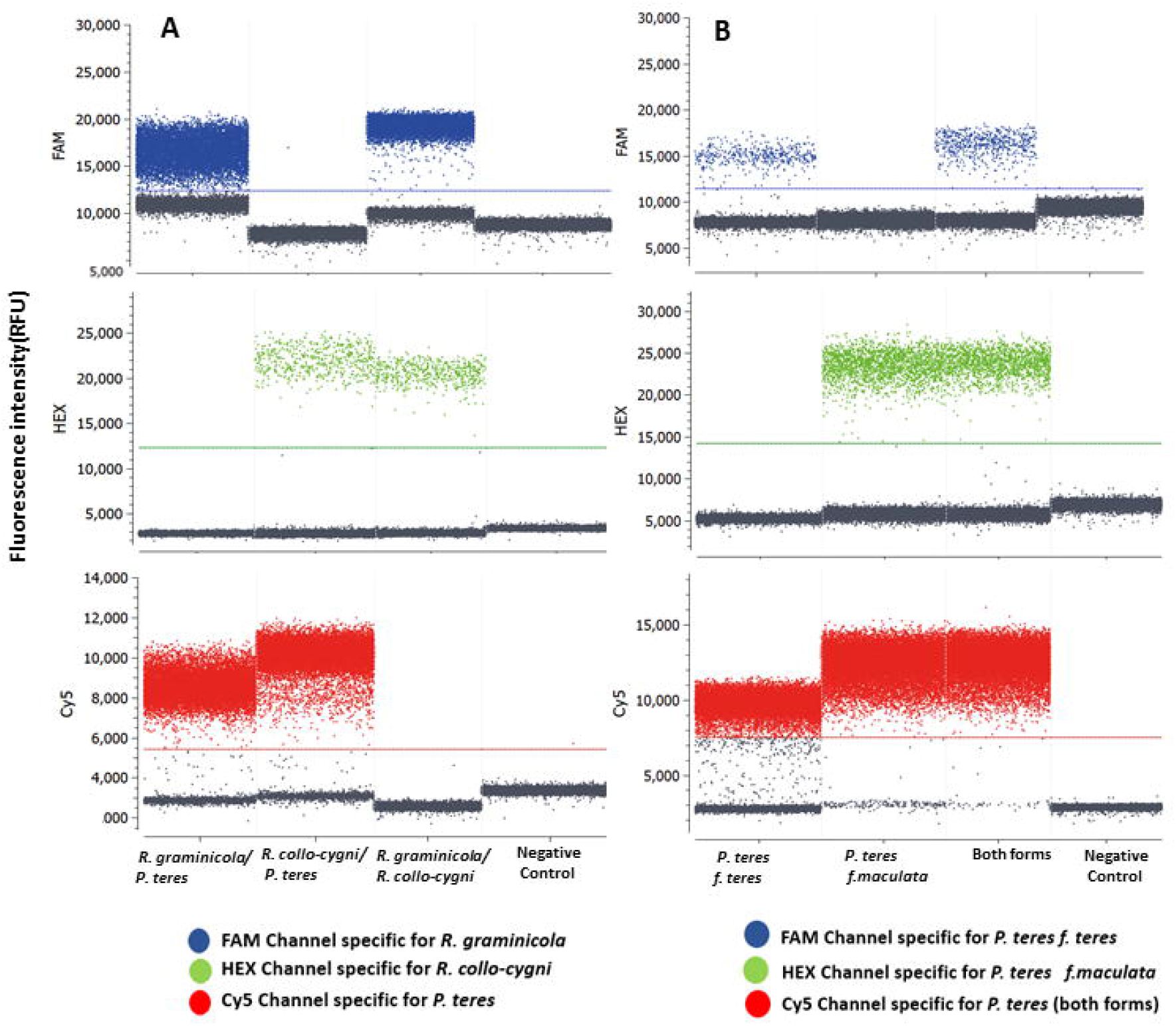
Digital droplet PCR assay, pathogen detected are indicated on x-axis and coloured horizontal line is indicating the threshold for the assay. Grey droplet population below the threshold line is negative droplets. Fluorescence intensity is indicated as relative fluorescence units (RFU)(A) Triplex assay for the detection of *R. graminicola, R collo-cygni* (cp/µl) and *P. teres* (cp/µl) (B) Form-specific assay for the detection of forms of *P. teres*, FAM channel detects *P. teres f. teres* while HEX channel is to detect *P. teres f. maculata. P. teres* ITS assay was the control (labelled with Cy5).

### 3.2 Linear dynamic range of the assay

To determine the accuracy of the assay, the linear dynamic range of the assay was ascertained by correlating the projected DNA copies of each target in a sample and copy numbers detected. For each probe-pathogen combination R^2^ values of >0.99 were calculated. Linear dynamic range of the assay also indicated the lower limit of detection of the assay, which is 0.00001 ng/µl (Figure 3). The linear dynamic range of the *P. teres* form specific assay was not determined as it was specifically designed to distinguish between forms following initial detection of *P. teres* using the triplex assay. Details of cp/µl for linear range is provided in Supplementary information (Linear dynamic range.xlsx).

**Figure 3.**
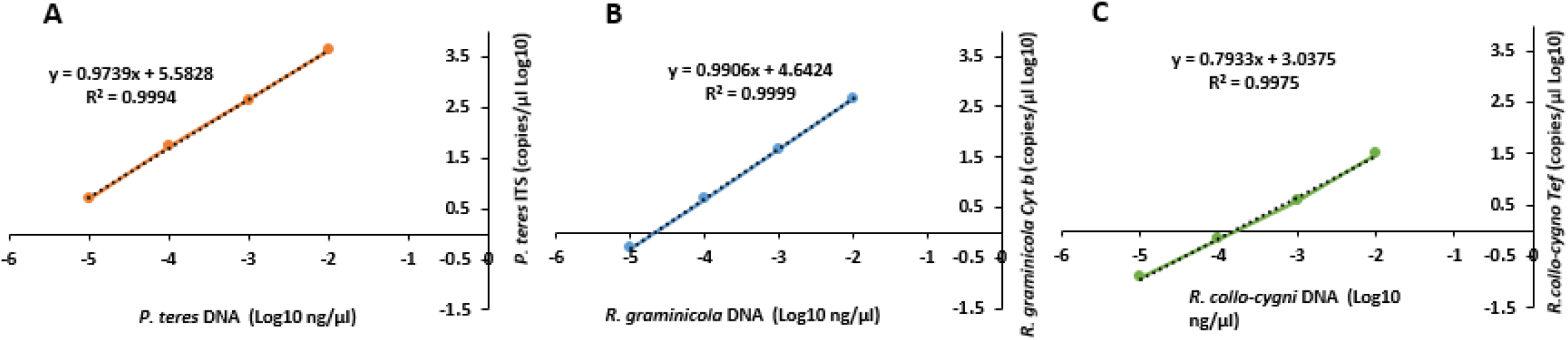
Linearity of the triplex assay for the detection of (A) *P. teres* (Internal transcribed Spacer (ITS) gene), (B) *R. graminicola* (Cytochrome b (Cyt b) gene) and (C) *R. collo-cygni* (Transcription elongation factor (tef) gene).

### 3.3 Specificity of the assay

Considering the potential application of our assays, for screening seeds and leaf samples from the field, specificity of both the assays was investigated against a range of fungi that may be found on or in barley seeds and leaves. No positive droplets were detected in any of the fluorescent channels for any of the fungi tested, indicating that the assays are highly specific (Figure 4A & B).

**Figure 4.**
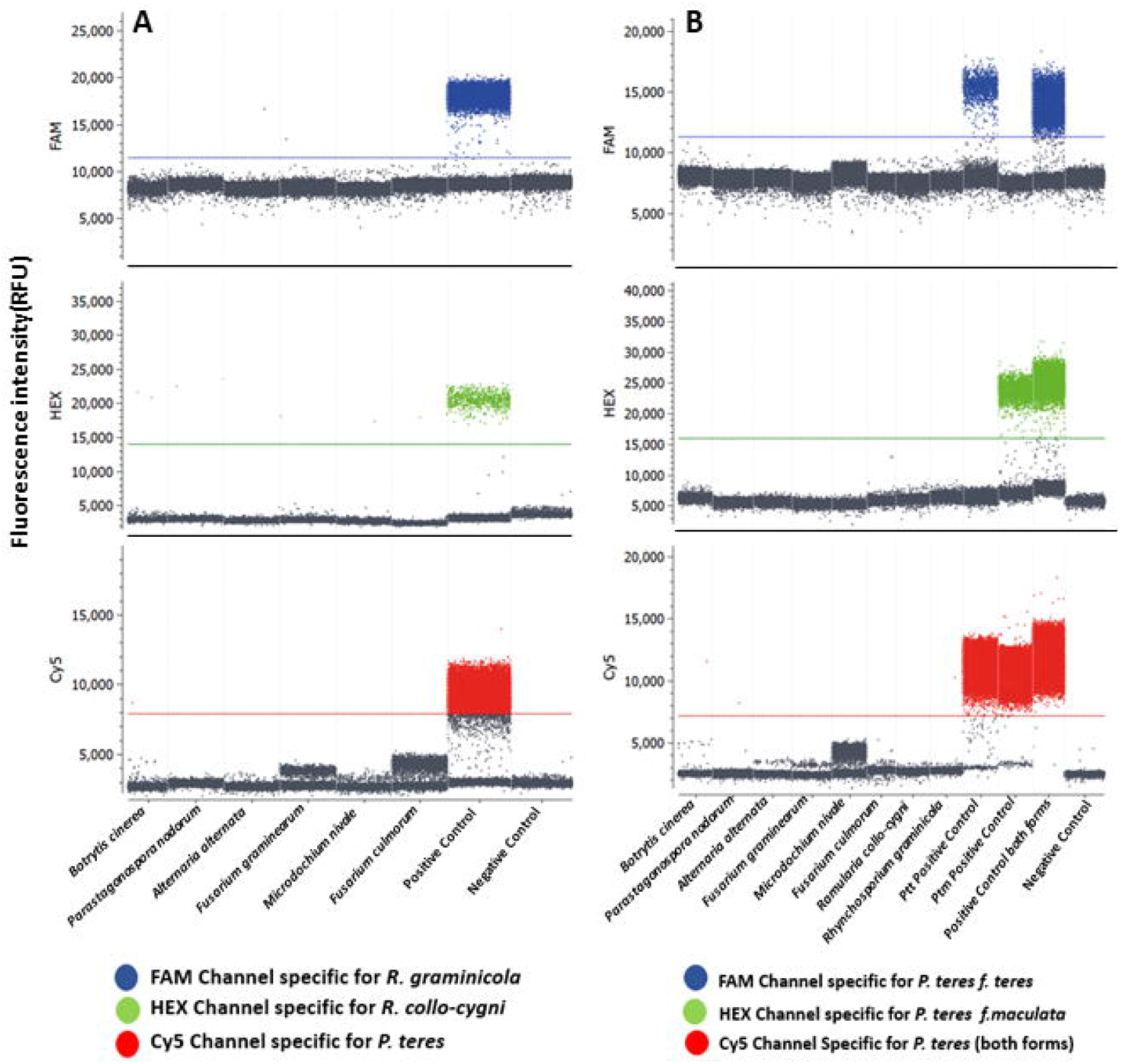
Specificity of the multiplex assays (A) Triplex assay for the detection of *R. collo-cygni*, R. *graminicola* and *P. teres* challenged with a number of barley pathogens (B) *P. teres* form-specific assay exposed to various barley pathogens.

### 3.4 Detection of barley pathogens in wheat

Wheat leaf DNA was also tested for the presence of *R. collo-cygni*. The assay showed a 469.7 cp/µl in sample 9, 4546 cp/µl and 1557 cp/µl in sample 11 and 13 respectively of *R. collo-cygni* DNA from symptomatic wheat leaves. However, wheat samples displayed positive droplets for the presence of *R. graminicola and P. teres* DNA in the samples (Figure 5). In order to rule out the *P. tritici-repentis* cross reactivity with the triplex assay, we performed the form-specific assay and we observed, 0.86 to 1.18 cp/µl of Ptt across two samples (Supplementary Figure 2).

**Figure 5.**
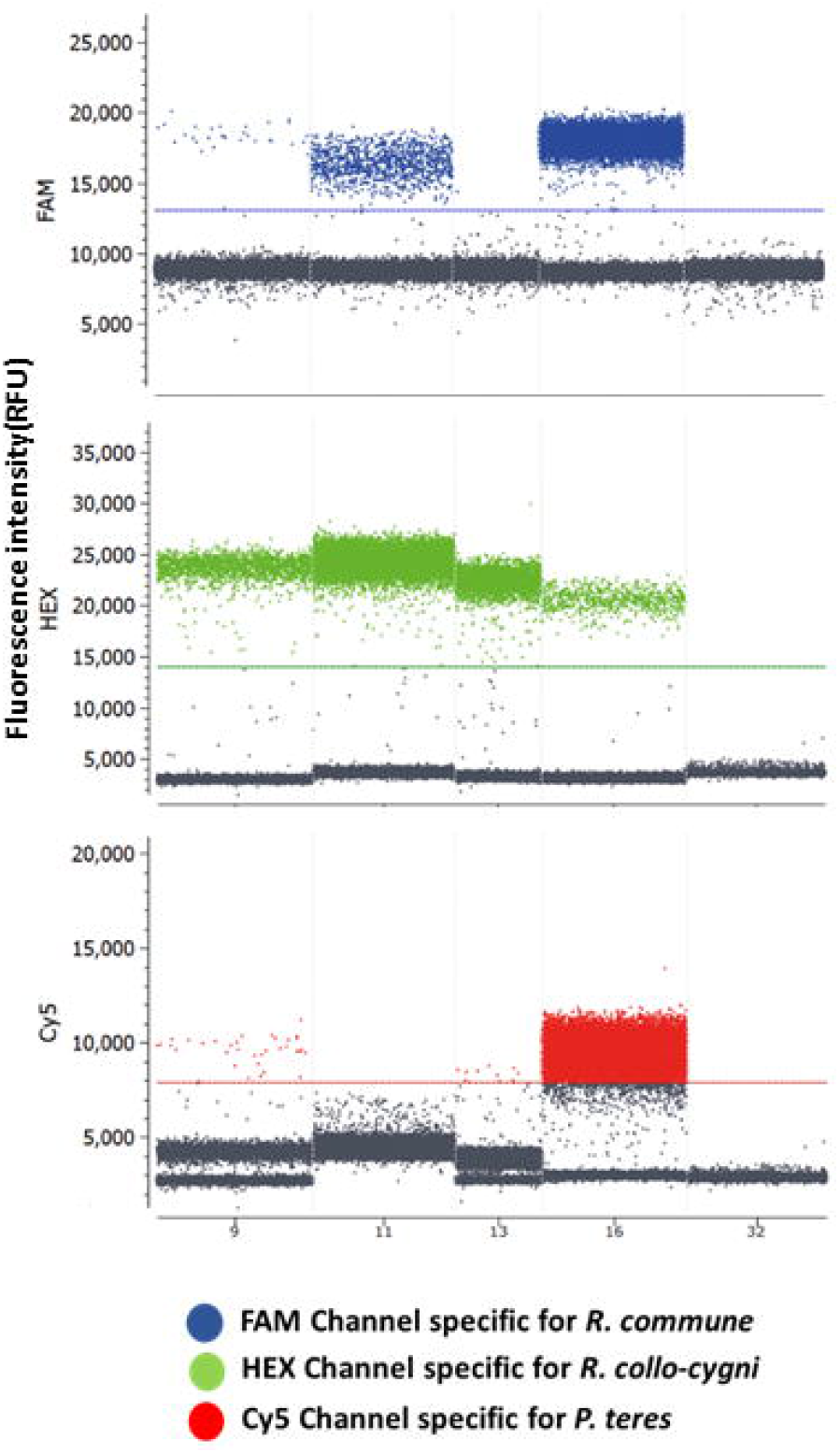
Detection of *R. collo-cygni, R. graminicola* and *P. teres* from wheat leaf DNA (sample 9, 11 and 13) collected from wheat volunteers in the barley field, sample 16 and 32 indicate positive and negative controls for the triplex assay. Droplet population in FAM and Cy5 channel indicate the possible cross contamination of the wheat leaves with *R. graminicola* and *P. teres* spores from the barley.

### 3.5 Detection of *R. collo-cygni, R. graminicola* and *P. teres* in barley seed samples

To determine the application of assays to test the presence of pathogens in naturally infected barley DNA samples, previously extracted from seeds of winter and spring barley from 2016 (Kildea et al. 2024) were tested. Only samples exhibiting cp/µl equal to or above the limits of detection, as identified above, were deemed positive for the respective pathogens. Of the 35 winter barley samples tested, 16 samples were positive for *R. collo-cygni*, two samples were positive for *R. graminicola*, while 19 were positive for *P. teres*. For the 32 spring barley samples, 15 were positive for *R. graminicola*, 25 samples were positive for *R. collo-cygni*, and 25 positive for *P. teres*. Nine of the spring and winter barley samples that were found to be positive for *P. teres* were tested to determine the form determination. None of the samples tested positive for the presence of *P. teres f. maculata*. In the case of winter barley, eight out of nine samples were positive for *P. teres f. maculata* and two samples were positive for *P. teres f. teres* as well as *P. teres f. maculata* (Supporting information seed testing data).

## 4 Discussion

Clean seeds are undeniably important for healthy crops and this is reliant on early detection of specific disease agents, which can further facilitate the execution of IPM based disease control strategies (Dussart et al. 2020; Tsedaley, 2015). The ddPCR assays described offer a specific, reliable and sensitive means of detecting and quantifying three of the most economically important barley pathogens in a single reaction. The end point nature of the digital PCR assay makes it ideal for detecting the different pathogens in complex plant materials, such as seeds (Liu et al. 2011).

To validate the efficacy of our assays and investigate the prevalence of *R. collo-cygni, R. graminicola* and *P. teres* in field samples, we screened a selection of historical barley seed samples collected from 2016 to 2018 and wheat leaves sampled from volunteer wheat plants in a winter barley field in 2023. Unsurprisingly, the barley seed samples were contaminated with variable levels of the three pathogens, with a proportion coinfected with all three pathogens. All three pathogens were detected in the winter wheat leaves, the latter result is unsurprising given the fact that the leaves tested were collected from wheat volunteers collected from a winter barley field. Whilst *R. collo-cygni* has previously been reported in wheat in Ireland (Zia et al. unpublished), the detection of *R. secalis* and *P. teres* in the sample may simply reflect the presence of both pathogens on the wheat leaf surface rather than colonisation of the leaf tissue. This highlights a potential limitation of the assays. As they are based on the detection of DNA of the respective pathogens, they cannot determine if the detected pathogen was actively colonising the plant tissue, or if the pathogen detected was even viable. However, it is important to note that these limitations are not restricted to the assays developed and described in this paper but are a drawback of all DNA-based detection methods (Ficetola et al. 2015). It is therefore critical that any positive detections are assessed in the context of detection (host) and the potential risk(s) posed by the pathogen. As mentioned previously it is more likely that the positive detections for *R. graminciola* and *P. teres* from the wheat volunteers are consequential to levels of both diseases in the surrounding barley crop. However, it is worth highlighting that in addition to *R. collo-cygni* being noted as able to colonise wheat, Ptt has been reported as causing disease on wheat in Hungary (Tóth et al. 2008), Russia (Mikhailova et al. 2010) and most recently in Brazil (Garozi et al. 2020). Further analysis of wheat grains and/or a wider wheat disease survey is warranted to determine if either pathogen is more widespread in wheat crops.

Those barley seed samples that presented positive for *P. teres* were subsequently tested to determine what form of *P. teres* was present. Amongst the spring barley samples tested only Ptt was present. For the winter barley seed samples both Ptt and Ptm were detected, with a number of samples positive for both. All samples with detectable levels of Ptm, irrespective of the presence of Ptt, were from the cultivar KWS Tower, which had by the late 2010s become highly susceptible to *P. teres*, and in particular Ptm (Recommendation list in supplementary information and author’s unpublished data). Positive detection of both Ptt and Ptm could indicate of a potential Ptt/Ptm hybrid, however based upon the number of droplets positive for both in the test samples this appears unlikely. The presence of markers for both forms does suggest that hybrids may arise, with recent reductions in fungicide sensitivity and efficacy in Western Australia associated with such a hybrid (Mair et al. 2020; Turo et al. 2021).

It is worth noting that within the multi-pathogen assay the region targeted for *P. teres* was ITS region, whilst in the form specific assay we targeted the Ptt form-specific region PttQ5 mRNA sequence and the Ptm form-specific region PtmQ11 mRNA sequence. The copy number for ITS in different fungi can vary considerably (Lofgren et al. 2019) and the copy number of PtmQ11 and PttQ5 are unknown in *P. teres*. As a number of samples that tested positive for *P. teres* in the multi-pathogen assay, were neither positive for either Ptt or Ptm suggests PtmQ11 and PttQ5 have a lower copy number compared to ITS region in *P. teres*. Therefore, ~ 15 cp/µl of *P. teres* sequence are required to confirm the presence of Ptt or Ptm from the samples, as observed in the case of historical seed samples. Moreover, high droplet population in *P. teres* ITS could potentially be the consequence of similarity between *P*.*teres* and *P. graminea* ITS regions (Supplementary Figure 1), therefore, we suggest further exploration of the historical seed samples.

As infected seeds are a likely source for the introduction of new pathogens and resistance /virulence mechanisms across the globe, including the three pathogens discussed, early detection and subsequent management in seed stocks will help minimise their spread (Stam et al. 2019). Therefore, molecular seed screening for pathogen levels and identification of the presence of fungicide resistance or virulence alleles should be a regular practice prior to seed transportation. This ddPCR assay can be an efficient tool for regular seed screening. Furthermore, establishment of pathogen threshold levels by deploying ddPCR, coupled with the rejection of seeds with high pathogen levels, can minimize the risk of future outbreaks. Seed is only one of the sources of these diseases, however prevention is the key to suppression.

## Supporting information

In the case of winter barley, eight out of nine samples were positive for P. teres f. maculata and two samples were positive for P. teres f. teres as

All fungal cultures used in the study (Supplementary Table 1)

## Acknowledgments

This study is part of BioCrop, a project funded by The Irish Department of, Agriculture, Food and The Marine under their Research Stimulus Fund (Ref: 2019PROG705). We would like to thank Dr. Diana Bucur (Teagasc, Oak Park) for fungal isolates for assay specificity testing.

## Data availability statement

The data that support the findings of this study are available on reasonable request from the corresponding author.

---

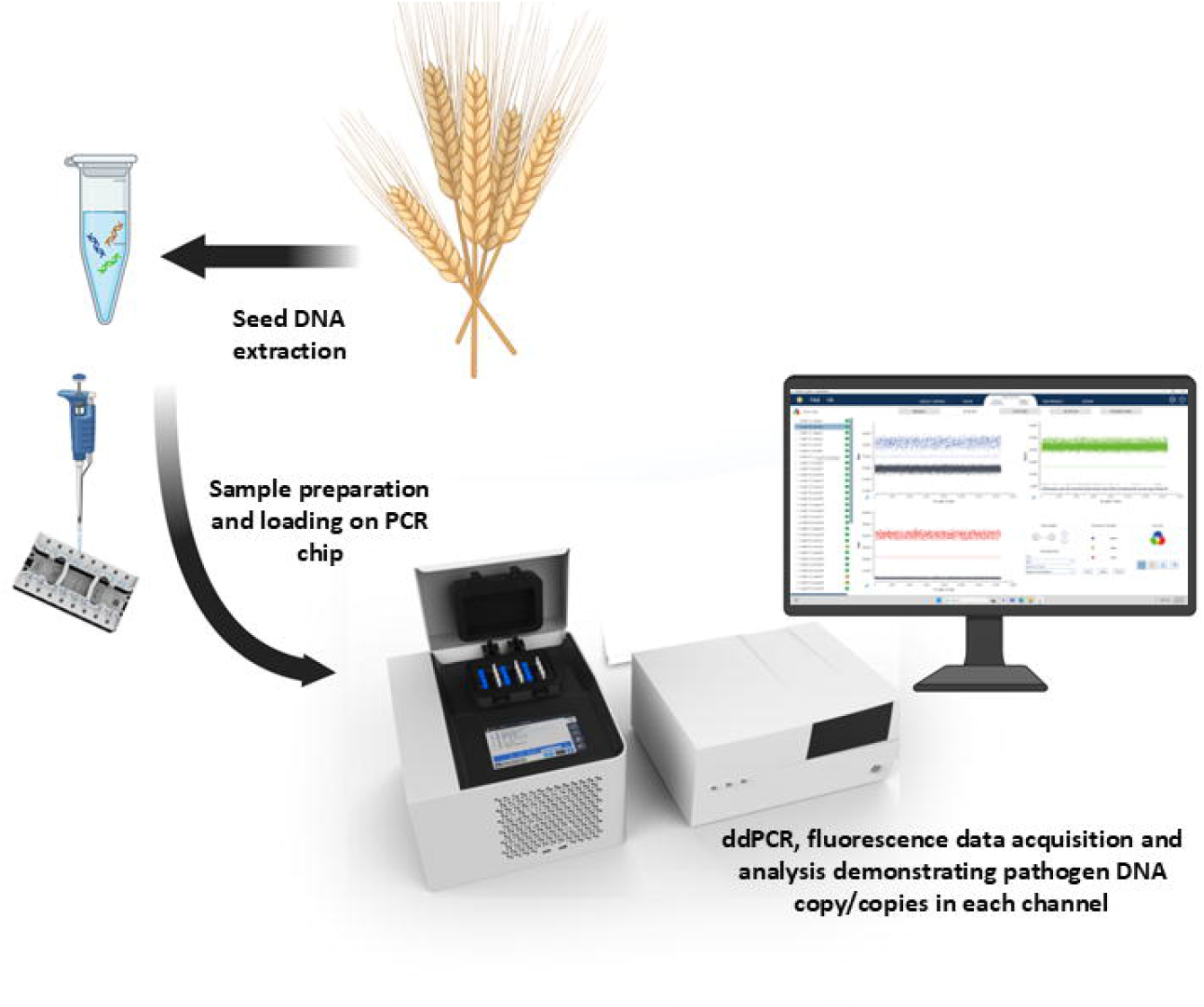

