## Supplementary material for "Barley disease screening: a multiplex digital droplet PCR approach for the detection of *Ramularia collo-cygni, Rhynchosporium graminicola* and *Pyrenophora teres*": All fungal cultures used in the study (Supplementary Table 1)

Figure S1

Routine PCR for *R. graminicola, P. teres* and *R.collo-cygni* primer optimization

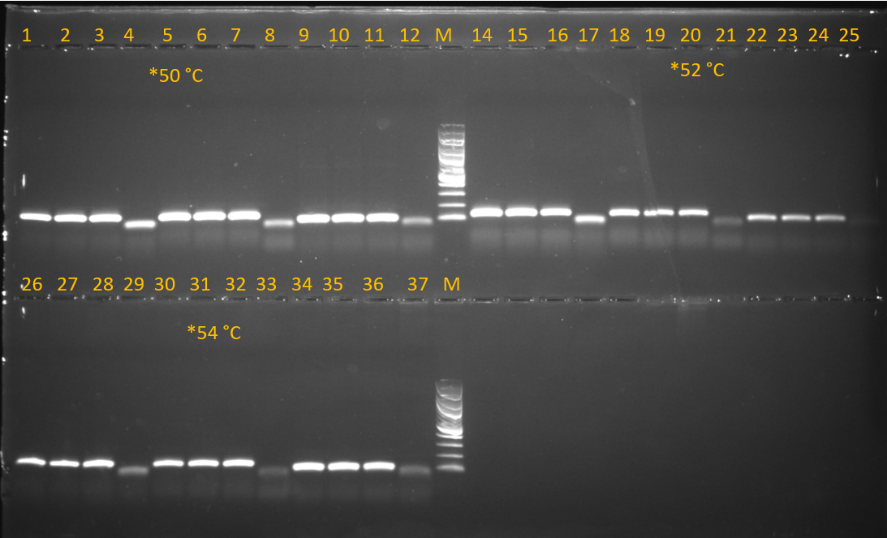
Routine PCR was conducted by choosing a range of annealing temperatures. PCR reaction mixture consisted of OneTaq Quick-Load 2X Master Mix (New England Biolabs, UK), 200nM of each forward and reverse primers were used and 5 µl of pure pathogen DNA without normalization was used. Three separate reactions mixtures were established for each pathogen.

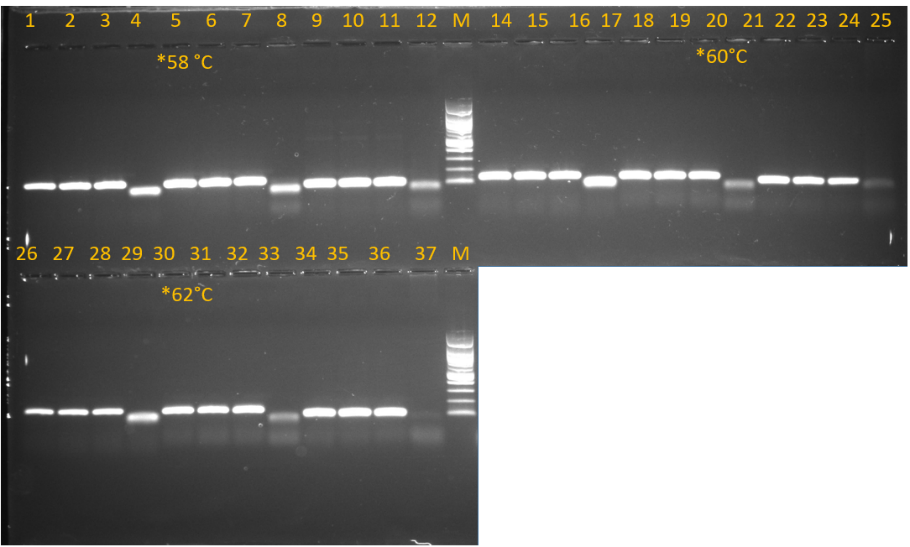

1-3, 14-16 & 26-28= *R. graminicola* 125 bp

4, 17 & 29= negative Control for *R. graminicola*

5-7, 18-20 & 30-32= *P. tere 123 bp*

8, 21 & 33= negative Control for *P.teres*

9-11, 22-24 & 34-36= *R. collo-cygni* 88 bp

12, 25 &37= negative Control for *R.collo-cygni*

M= 100 bp DNA ladder (Thermo Scientific™.US)

*50 °C, 52 °C 54 °C, 58°C 60 °C and 62 °C were six tested annealing temperatures, however, We tested until 62°C by PCR however, as there was no observable drop-off in performance at the highest temperature we selected to start with 64°C in ddPCR and subsequently kept it as a final annealing temperature.

Supplementary Table 1

| No. | Isolates | DNA concentration used for ddPCR | Isolate code | Host species | Origin |
| --- | --- | --- | --- | --- | --- |
| *1* | *Botrytis cinerea* | 20 ng/µl | Bc_1(2) | Faba beans | Oak Park, Ireland |
| *2* | *Alternaria alternata* | Presence confirmed with PCR^*^ | NA | Wheat | Ireland |
| *3* | *Fusarium graminearum* | Presence confirmed with PCR | NA | Wheat | Ireland |
| *4* | *Microdochium nivale* | Presence confirmed with PCR | NA | Wheat | Ireland |
| *5* | *Parastagonospora* *nodorum* | 2.28ng/µl | Pnod_123 | Wheat | Ireland |
| *6* | *Fusarium* *culmorum* | Presence confirmed with PCR | NA | Wheat | Ireland |
| *7* | *R. collo-cygni* | 0.1 ng/µl | R4.9.2 | Barley | Oak Park, Ireland |
| *8* | *R. graminicola* | 0.1 ng/µl | R18.23.4.5 | Barley | Oak Park, Ireland |
| *9* | *P.teres f. teres* | 0.1 ng/µl | Ptt_123 | Barley | Oak Park, Ireland |
| *10* | *P.teres f. maculata* | 0.1 ng/µl | Ptm_123 | Barley |  |
|  | *R. secalis* | 1.05 ng/µl | Rsc_123 | Rye | Oak Park, Ireland |

Supplementary Table 1

Isolates utilized in the study

***P. teres* target sequence with primers and probe placement**

**>OM471874.1**

Forward Primer

3’TTCGAGCGTCATTTGTACCCTCAAGCTTTGCTTGGTGTTGGGCGTCTTTTGTCTCTCCCCCGAGACTCGCCTTAAAAACATTGGCAGCCGGCCTACTGGTTTCGGAGCGCAGCACATTATTTG 5’

Reverse Primer Primer

Hydrolysis probe

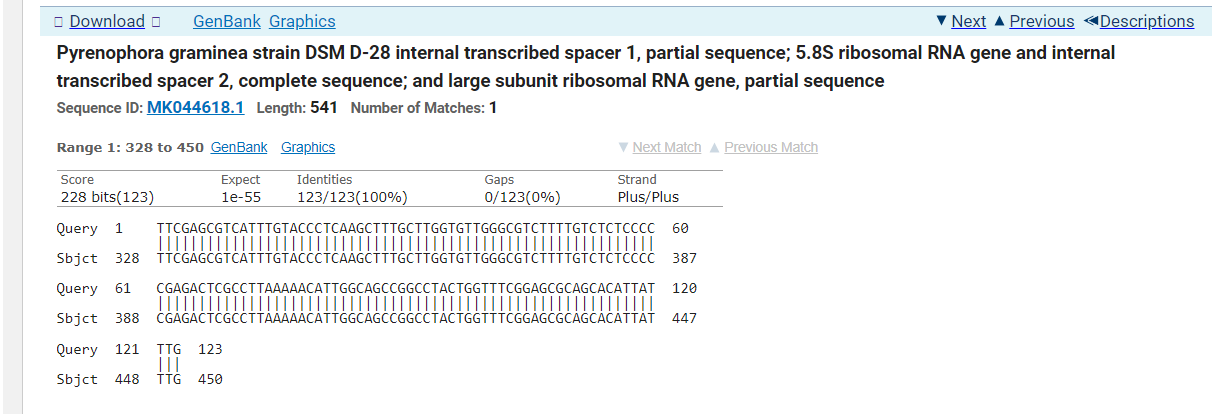

**Supplementary Figure 1**

Supplementary Figure 1: BLAST search indicating sequence similarity of *P. teres and P. graminea*

**Supplementary Figure 2**

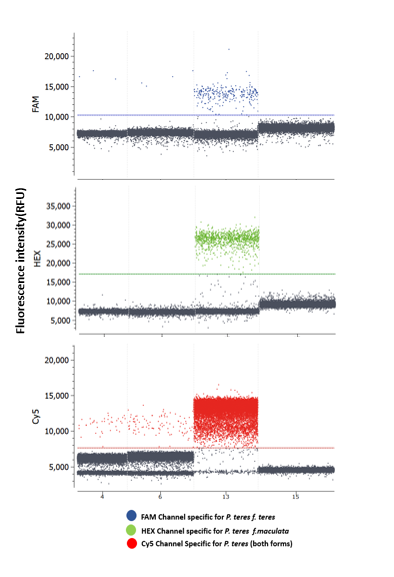

**Supplementary Figure 2:** Wheat samples positive for*P.teres* positive samples were tested with form-specific assay and we found few droplets in *P.teres f.teres* channel as shown in figure below, however, the copy number for *P.teres f.teres* is below detection limits for the Ptt assay. Figure provided below; where 4 and 6 are wheat samples while 13 and 15 are positive and negative controls respectively
